# Network for Knowledge Organization (NEKO): an AI knowledge mining workflow for synthetic biology research

**DOI:** 10.1101/2024.06.27.601082

**Authors:** Zhengyang Xiao, Himadri B. Pakrasi, Yixin Chen, Yinjie J. Tang

## Abstract

Large language models (LLMs) can complete general scientific question-and-answer, yet they are constrained by their pretraining cut-off dates and lack the ability to provide specific, cited scientific knowledge. Here, we introduce <u>Ne</u>twork for <u>K</u>nowledge <u>O</u>rganization (NEKO), a workflow that uses LLM Qwen to extract knowledge through scientific literature text mining. When user inputs a keyword of interest, NEKO can generate knowledge graphs and comprehensive summaries from PubMed search. NEKO has immediate applications in daily academic tasks such as education of young scientists, literature review, paper writing, experiment planning/troubleshooting, and new hypothesis generation. We exemplified this workflow’s applicability through several case studies on yeast fermentation and cyanobacterial biorefinery. NEKO’s output is more informative, specific, and actionable than GPT-4’s zero-shot Q&A. NEKO offers flexible, lightweight local deployment options. NEKO democratizes artificial intelligence (AI) tools, making scientific foundation model more accessible to researchers without excessive computational power.

## Introduction

Biomanufacturing has a potential US market of over 30 billion dollars annually (2023 Government Accountability Office report). However, a primary challenge that needs urging solutions is the high cost to develop cellular factories that meet commercially relevant performance. Biological systems are complex and there are many important levers (e.g. genetic regulations, enzyme functions, cellular metabolism, and extracellular conditions) that need to be analyzed and tuned to engineer a desired phenotype (i.e., design-build-test-learn cycles for bioprocess development) ^1^. Therefore, a holistic knowledgebase for bioprocess development is essential. Currently, vast amount of synthetic biology and biomanufacturing literatures have been published. A PubMed search of “synthetic biology or metabolic engineering” queries generated ∼125,000 publications, which offers wealthy bioinformation. The pressing need for efficient knowledge integration has coincided with the advent of large language models (LLMs) ^2,3^, which now facilitate rapid text information processing. Notably, recent advances in retrieval-augmented generation (RAG) have made LLMs powerful tools for information processing and knowledge mining from text ^4-6^. While general-purpose pretrained LLMs can provide general answers to scientific inquiries ^7^, encapsulating infinite knowledge within LLM’s finite parameter space remains an inherent challenge ^8,9^. Therefore, LLM needs a database to store factual knowledge. Knowledge graphs have emerged as a promising solution, offering an intuitive representation and knowledge synthesis that is interpretable by both LLMs and humans ^10,11^. The synergy between LLM and knowledge graph construction can help scientists collect key information and make informed decisions on research.

In this study, we introduce NEKO, a versatile knowledge mining workflow exemplified through case studies, demonstrating its broad applicability in data collections and information distillations from diverse reliable sources (**Fig. 1)**. First, NEKO helps scholars to quickly inquire into a research topic by combining, distillate and organize information from multiple articles. Second, NEKO can solve logic problems and generate hypothesis to be tested by connecting key concepts and synthesizing holistic knowledge from closely relevant articles. Third, it can help experiment planning and troubleshooting by presenting all past considerations in a systematic way. Therefore, NEKO can not only accelerate the learning and training process of synthetic biology researchers, but also assist them to write literature review, prepare research proposal, and optimize experimental design. NEKO is compatible with any instruction-following LLM, including proprietary models like GPT-4 (Open AI) ^2^ and open-source alternatives like Qwen (Alibaba) ^3^. Recently, the concept of foundation model ^12,13^ becomes a new research interest due to its specialized knowledge in various fields. NEKO is a lightweight application that anyone can construct synthetic biology foundation model according to specific needs. Developed using beginner-friendly Python code, the entire workflow is readily accessible via GitHub.

**Figure 1.**
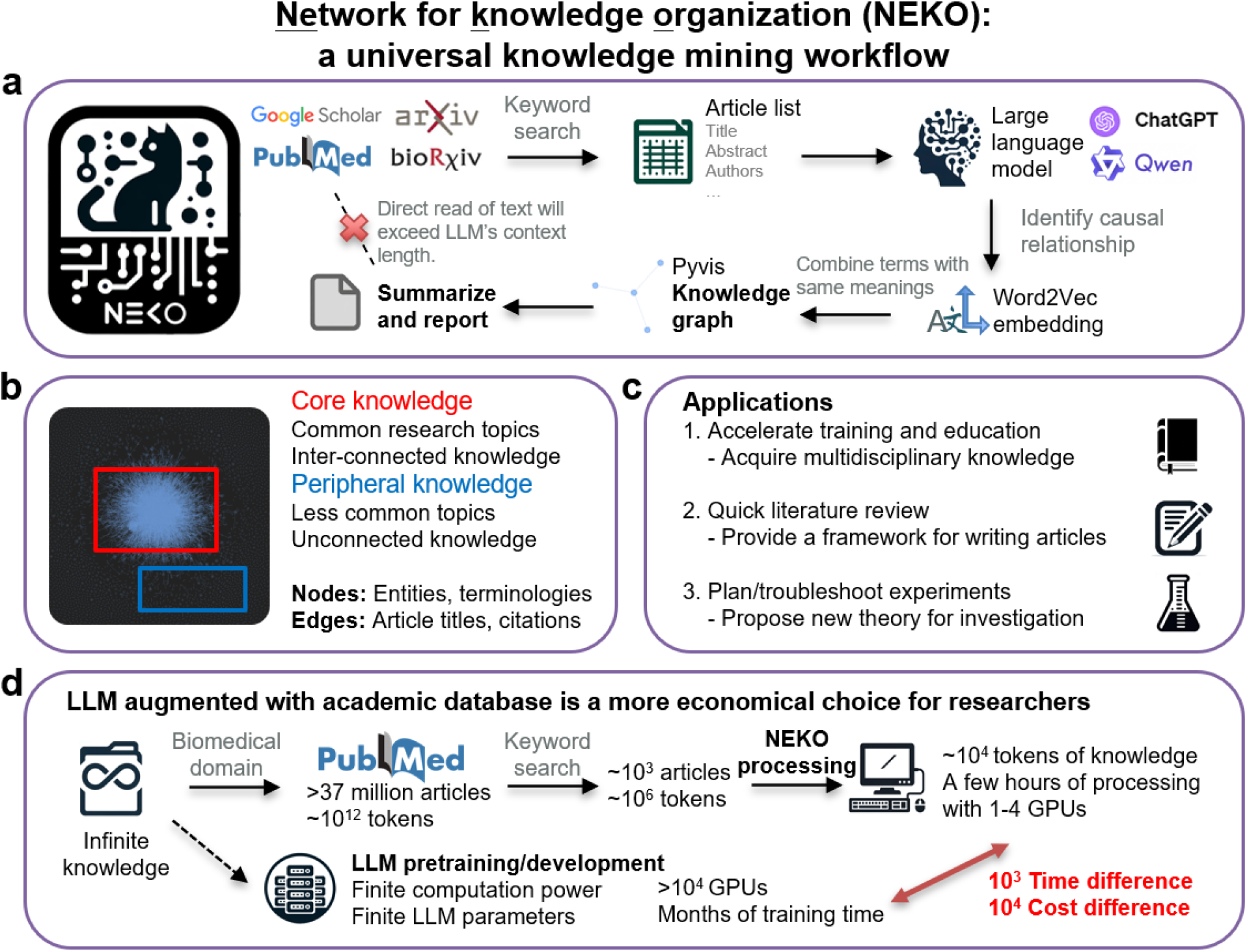
NEKO: a universal knowledge mining workflow. (a) NEKO’s logo and workflow illustration. (b) Knowledge graph has two regions, core and peripheral. (c) Applications in academic tasks. (d) LLM augmented with academic database is a more economical choice for researchers.

## Results

NEKO (“cat” in Japanese, **Figure 1a)** is a generative AI tool which can be a companion for synthetic biology researchers. Using *Y. lipolytica* PubMed abstract knowledge mining as a demonstration (**Figure 1b**), we plotted all information on a single knowledge graph. We identified two main regions: the core knowledge region at the center, which encapsulates common research topics and interconnected knowledge from various articles, critical for tasks such as process optimization and scale-up in bioproduction applications. The peripheral knowledge region contains fewer common topics. Researchers are encouraged to integrate this peripheral knowledge with the core by conducting experiments. We demonstrated NEKO’s applicability in academic tasks through several case studies (**Figure 1C**). This method of augmenting LLM with existing academic infrastructure is a more economical choice. Users can locally deploy NEKO and quantized LLM even on consumer-level desktops, with as little as 24 GB GPU RAM. Compared to pretraining science LLM, NEKO saves tremendous time and monetary costs (**Figure 1d**).

We choose Qwen as an example LLM in this study due to its exceptional Retrieval Augmented Generation (RAG) ability. With 14B parameters, Qwen can achieve comparable RAG score as GPT-4 ^14^. The use local LLM is more economical and can bypass the request rate limitation of OpenAI API. Interestingly, during our practical deployment of NEKO, we found that Qwen1.5 is as good as GPT-4 in terms of instruction following, scientific reasoning, and the comprehensiveness of responses. To date, there was few Qwen application journal articles, leading us to conclude that Qwen1.5 is an underappreciated yet reliable LLM for practical applications.

### Case study 1: Knowledge acquisition and distillation on β-Carotene Production in *Y. lipolytica*

NEKO can help researchers quickly acquire up-to-date knowledge about one research topic. To illustrate, NEKO rapidly read more than 200 research articles in PDF files and produce a knowledge base for the oleaginous yeast *Y. lipolytica*. When searching for β-carotene production in *Y. lipolytica*, NEKO’s analysis pinpointed relevant nodes, with the summary presented in **Figure 2**. NEKO also produced knowledge graph illustrating the connections between knowledge entities (**Figure S1**). Users can click one line connecting two nodes and view the title of the article containing the knowledge connection. Compared to ChatGPT-4 zero-shot Q&A, NEKO gave 200% more gene targets, 200% more strain engineering strategies, and 57% more bioprocess cultivation conditions, with knowledge from 37 reliable peer-reviewed sources (**Table 1**). We also did a negative control by using base LLM Qwen1.5 to write a summary about β-carotene production in *Y. lipolytica* (**Figure S2**). Qwen1.5 only gave general optimization suggestions like GPT-4 did. This demonstrates that without NEKO knowledge mining workflow, the LLM does not has specific knowledge about β-carotene production. NEKO augments the LLM by providing a light-weight knowledge database.

**Figure 2.**
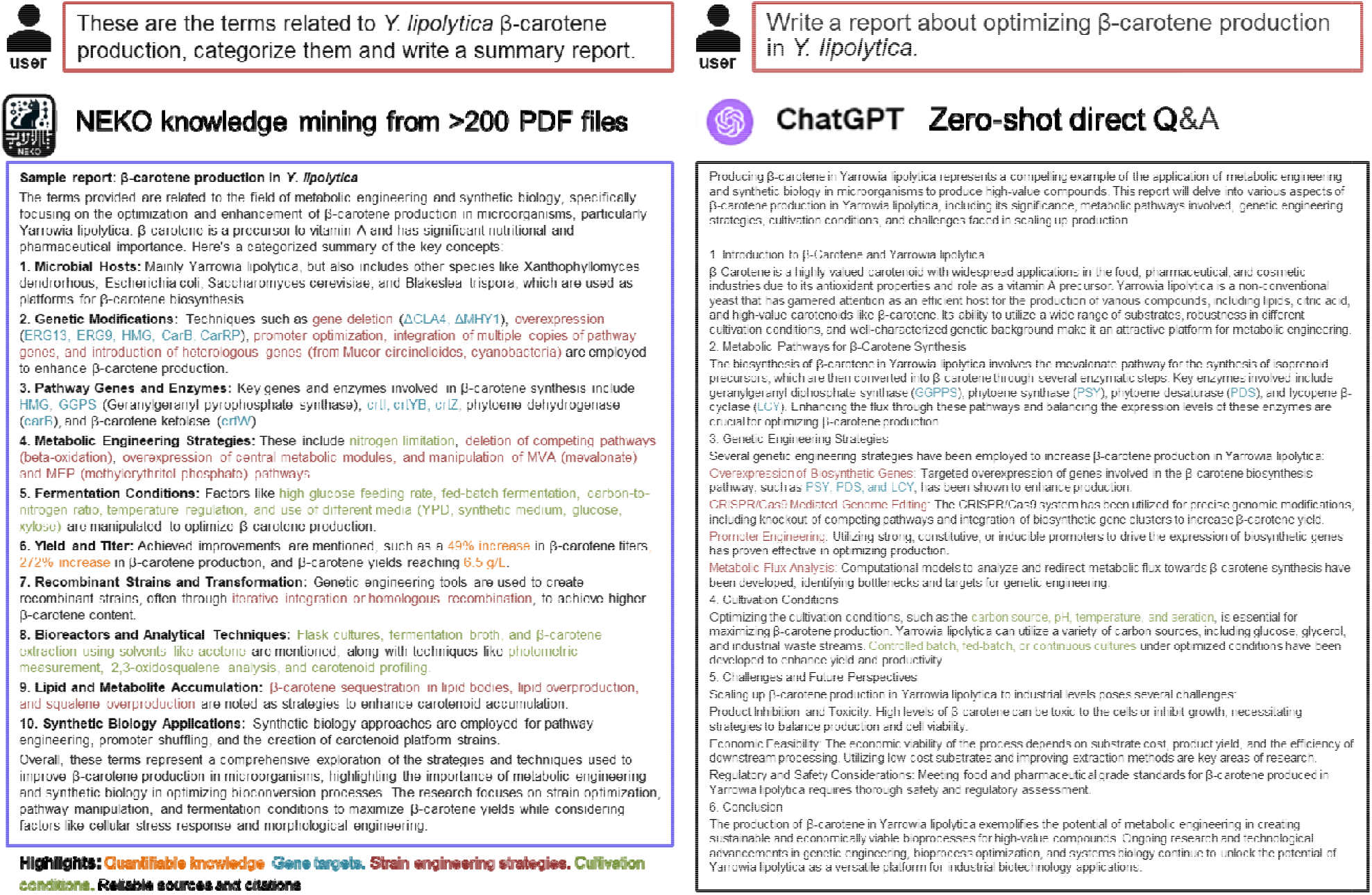
Compare NEKO and ChatGPT zero-shot Q&A. The topic is to write a summary of production of β-carotene in *Y. lipolytica*.

**Table 1:** Compare NEKO and ChatGPT zero-shot Q&A.

|  | NEKO workflow | GPT-4 zero-shot Q&A |
| --- | --- | --- |
| Quantifiable knowledge | 3 | 0 |
| Gene targets | 12 | 4 |
| Strain engineering strategies | 12 | 4 |
| Cultivation conditions | 11 | 7 |
| Reliable sources and citations | 37 | 0 |

Another information-intensive academic task is experiment planning and troubleshooting. We illustrate this through NEKO’s ability to synthesize experiment procedures and actionable insights. For example, when aiming to genetically engineer *Y. lipolytica*, user needs to obtain the procedures for strain transformation. NEKO provided a detailed, step-by-step methodological guidance (**Figure S3**), including specific DNA amount used (2 μg), procedure name (LiAc or electroporation), selection markers (hygromycin resistance or ura3d4 defective allele), etc. Take another example, assume the user is having trouble expressing GGPPS, an essential enzyme in β-carotene synthesis pathway. NEKO can search for information regarding to GGPPS and compile a report with suggestions to troubleshoot this scenario (**Figure S4**). We noticed that GPT-4’s suggestions were heavily templated. When given another β-carotene production gene CarRP (**Figure S5**), GPT-4’s suggestions were very similar to GGPPS’s case. Contrasting NEKO’s outputs with GPT-4’s templated responses highlighted NEKO’s ability to generate pertinent suggestions based on mined knowledge.

### Case study 2: Literature review of non-model species *Rhodosporidium toruloides*

Literature review is one of the time-consuming academic tasks, and we demonstrated NEKO’s quick application in review paper writing in this case study. *Rhodosporidium toruloides* is gaining research attention recently for its high lipid content and native carotenoid production. We applied NEKO to *R. toruloides* article abstracts (total 392 articles) on PubMed. By visualizing the knowledge graph and comparing with *Y. lipolytica* (**Figure S6**), *R. toruloides* literature is still at its early stage of research. The knowledge from studies is disconnected from each other. After LLM summarization **(Figure 3 and Figure S7)**, NEKO identified trending research areas, including genetic engineering, metabolic pathway, adaptive evolution experiments, and biochemical production. Moreover, NEKO enables users to refine their searches by specifying target products. For instance, focusing on lipid production yielded a detailed breakdown of the research topic (**Figure S8**). NEKO excels at providing details like strain engineering gene targets and fermentation conditions. Overall, NEKO provides a framework for users to organize the knowledge and structure their review paper writing. By further inspecting the article list and entity list, users can produce rich and precise literature study.

**Figure 3.**
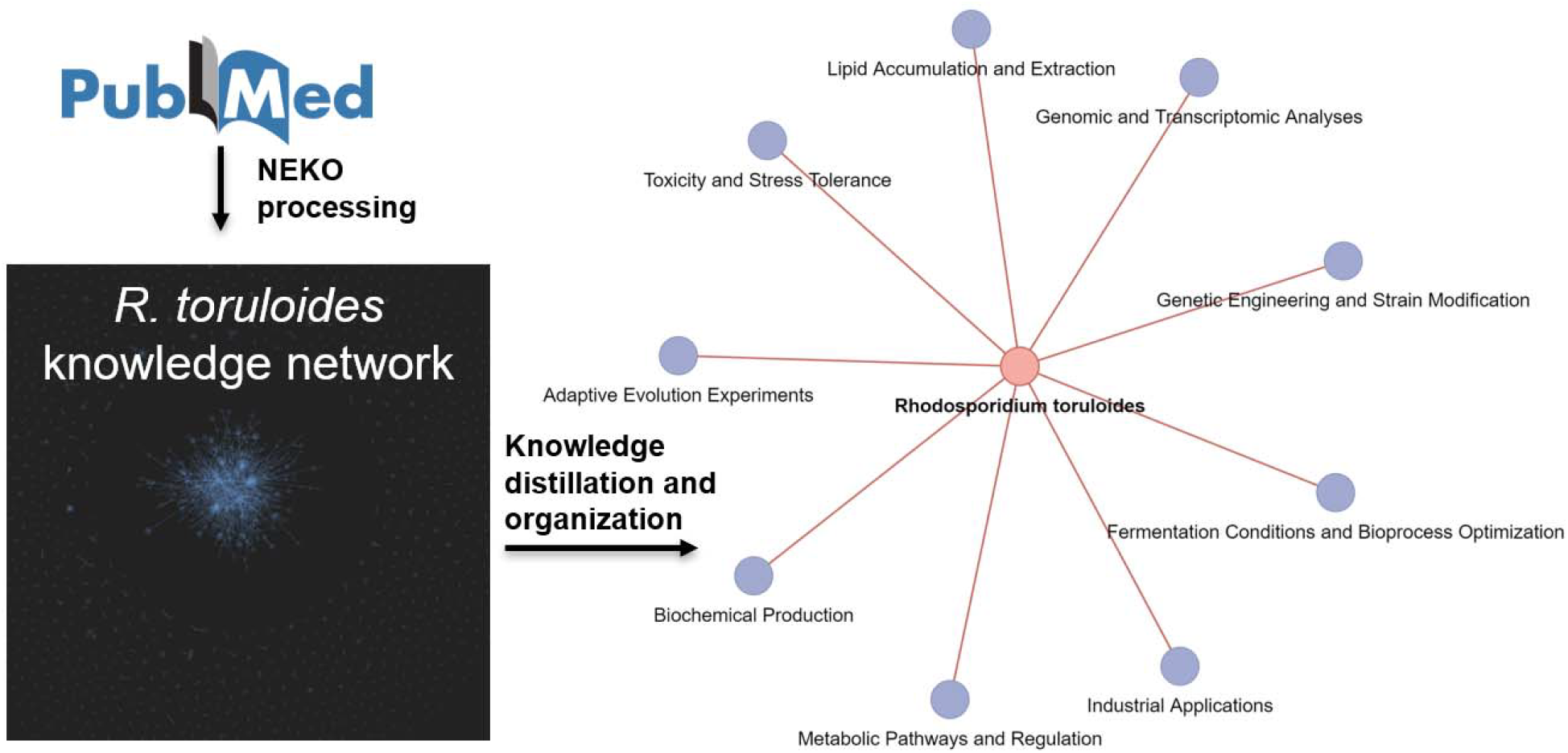
NEKO processes *R. toruloides* publications on PubMed and produced a quick research topics review.

### Case 3: Hypotheses generation and new theory for exploration

Being able to summarize literature information is part of a competent synthetic biology foundation model. In this case study, we demonstrate that LLM can do inference and propose new hypotheses given relevant concepts. In essence, relevant concepts were “prompts” for LLM to generate new experiment plan. For example, we are interested in engineering a nitrogen-fixing cyanobacteria *Anabaena* sp. to produce guanidine and urea as bio-fertilizers. We combined relevant concepts of *Anabaena* sp. nitrogen fixation and common bio-fertilizer formulations. The LLM Qwen1.5-72B can do inference and write a brief research proposal and experiment plan (**Figure 4, Figure S9**). Interestingly, same as our research plan, this workflow correctly identified gene targets as nitrogenase, arginine decarboxylase (ethylene-forming enzyme), and urease. Compared to GPT-4 zero-shot Q&A in **Figure S10**, NEKO pointed out that urea and guanidine are more stable than ammonia, and they are more suitable for long term storage and transportation. The proposed experiments are detailed, incorporating tools like CRISPR-Cas9, metabolic engineering, and dynamic expression control systems. In terms of project deliverables, NEKO’s report had a thorough plan that includes *in vitro* and *in vivo* characterization, field trials, and an economic/environmental evaluation, offering a more holistic project than GPT-4’s zero-shot Q&A.

**Figure 4.**
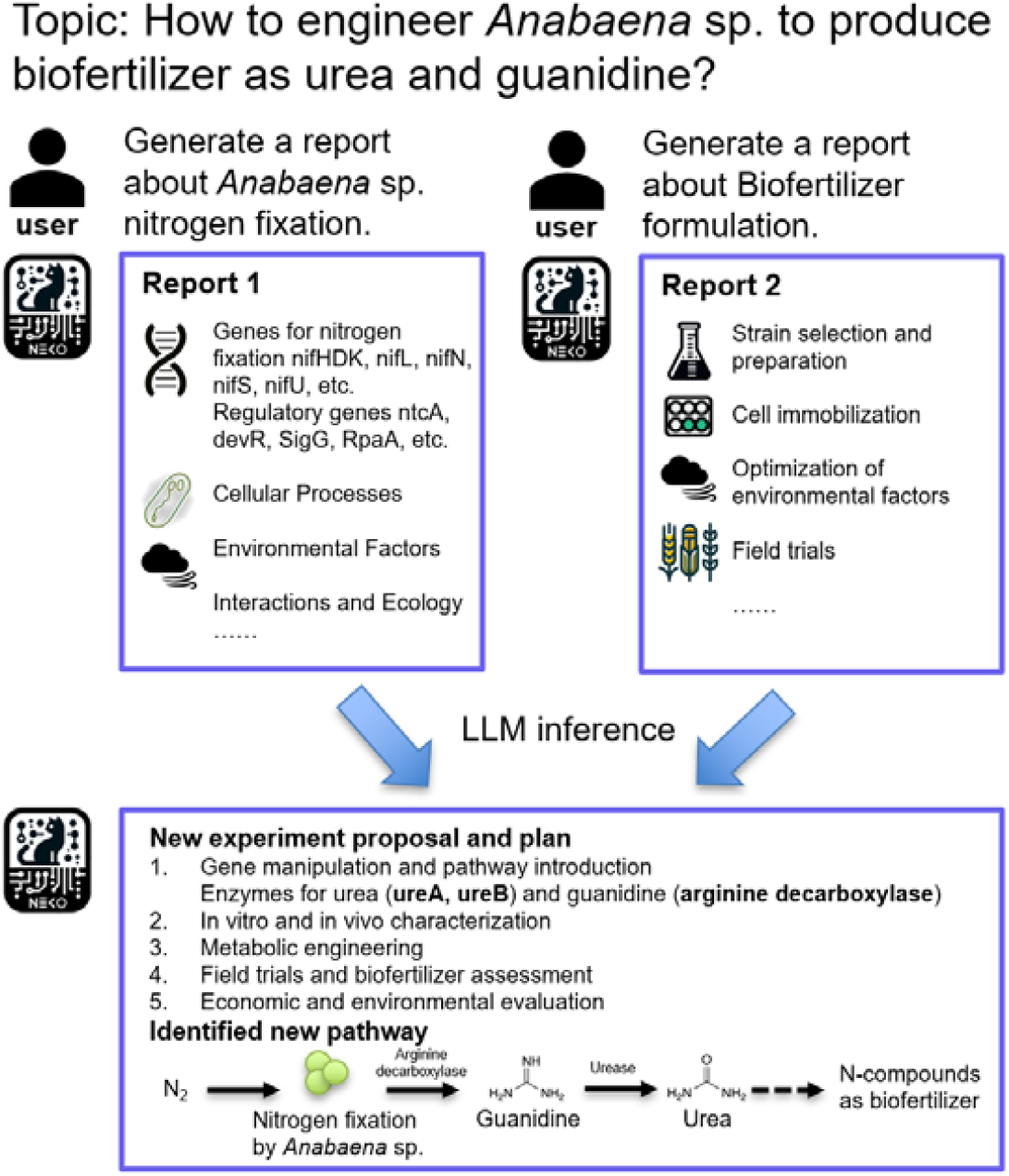
Based on the knowledge mining of NEKO, LLM can perform inference and give new research plan about new compounds production pathway.

**Figure 5.**
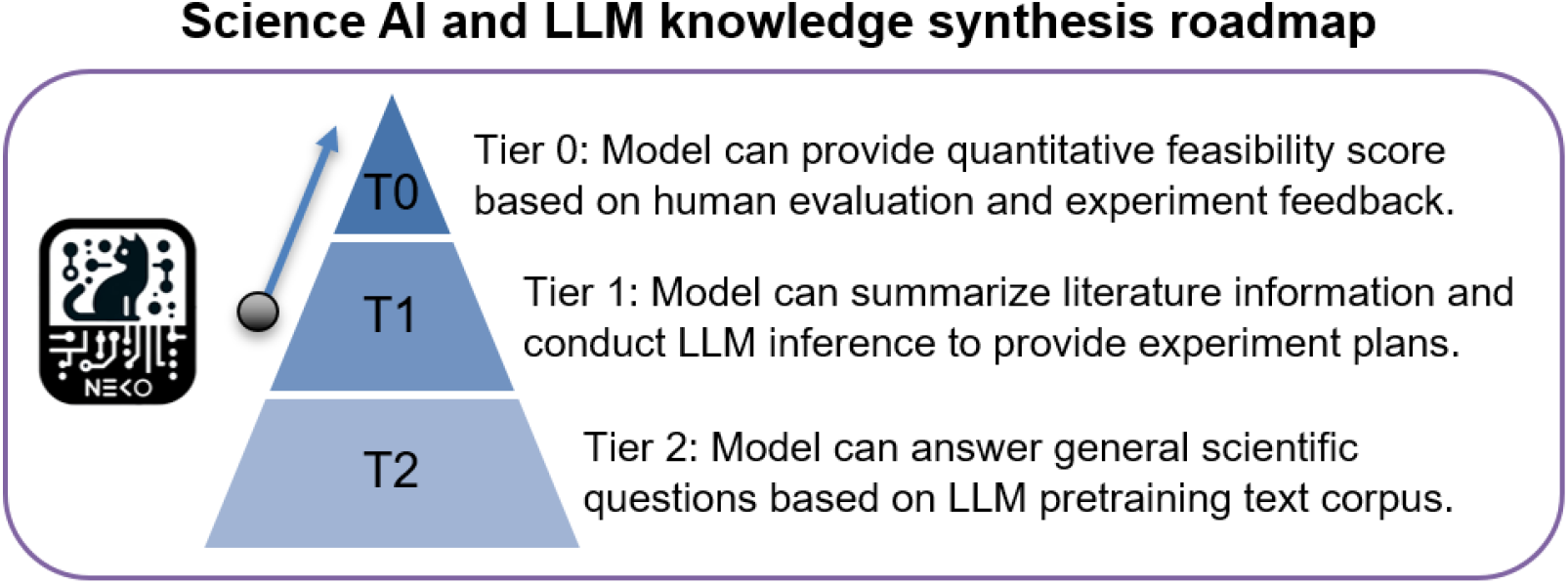
Proposed science AI and LLM knowledge synthesis roadmap.

## Discussion

NEKO can compile massive literature reports, fill knowledge gap, remove redundant data, and connect information streams, which can be used to collect both features and targets from literature for developing standardized datasets ^15^. NEKO can be widely used for Synthetic biology research. It distinguishes itself from other knowledge-based Q&A platforms by overcoming the context length limitations of LLMs without directly read article texts. All NLP tools used in this study are off-the-shelf, without the need for pretraining. NEKO is compatible with any LLM, although the effectiveness is influenced by the LLM’s ability to follow instructions and its RAG capabilities. Generally, LLMs with a higher number of parameters tend to perform better in identifying causal relationships and generating logical summary reports. When using Qwen1.5-72B running on 4*A100 GPUs, NEKO can process about 300 PubMed abstracts in 1 hour. It seems that GPU memory (GPU RAM) and CPU computation are the limiting factors. If using GPT-4, user’s API request rate is subject to OpenAI policies. Our preliminary findings indicate that NEKO typically offers more information compared to GPT-4. However, users are advised to verify important information. We also applied this workflow to other disciplines such as computer science. By modifying the knowledge mining prompt, NEKO was capable of produce a mini review of recent knowledge retrieval studies on arXiv (**Figure S11**). In general, NEKO works with any online article database with API services. Users need to modify the knowledge mining prompt based on their specific needs. We will constantly update our GitHub repository for more usage examples.

As academic research grows in complexity and increasingly spans multiple disciplines, researchers are investing substantial time in literature review and knowledge acquisition. NEKO addresses this challenge by streamlining literature study tasks, enabling knowledge synthesis from literature ^16,17^. This trend towards continuously interacting research with LLM represents a new common. We envision a roadmap for three tiers of science AI and LLM knowledge synthesis (**Figure 7**). Currently most pretrained LLMs are at tier 2, and they can answer general scientific questions based on pretraining text corpus. NEKO is at tier 1. It can output summary reports based on literature search and give reliable cited knowledge. To some extent, it can help guide experiments in the physical world. It serves as a “human prompter” by brainstorming potential experiment plans. The top tier 0 is one of the ultimate development goals for science AI and foundation model, which can provide quantitative feasibility score based on human evaluation and experiment feedback. However, there are challenges associated with this goal. First, data generated by experiments are multi-modal, and it is difficult to integrate multi-omics datasets by text LLM only ^18,19^. Multi-modal LLM should be able to process images, videos, and data plots, minimizing manual efforts for data curation. Second, the inherent variability of experiment ^20^ requires the establishment of standardized workflow to facilitate effective feedback loops between practical research and LLM inferences. Third, biological systems are chaotic and subject to the influence of experiment initial conditions ^21-23^, so the notion of creating an omniscient AI Laplace’s demon is not feasible. A pragmatic approach acknowledges these limitations, focusing on the integration of AI within a defined set of constraints and assumptions. The key is to integrate knowledge mining workflow with physical world, and NEKO provides a lightweight deployment example for integrating LLM with routine education and academic tasks. In the future, we will test this knowledge mining workflow coupled with experiment feedback, facilitating design-build-test-learn (DBTL) cycles and offering a foundation model for synthetic biology applications.

## Methods

### Code availability

The codes used in this study are deposited in this GitHub repository: https://github.com/xiao-zhengyang/NEKO

### Web-based literature search

This workflow starts with online literature search. This step is compatible with any web-based databases with application programming interface (API) services. In this study, we used PubMed and arXiv as examples. User of NEKO input a search keyword into corresponding API, and the API would return the article title and abstract. The search result was saved in an article list excel file.

This workflow also applies to PDF files. After downloading research articles in PDF, the files were read and divided into 1000 words segments. Then these texts were passed to the next step.

### LLM text processing

LLMs were used to process the text from API search or from downloaded PDFs. LLMs with strong retrieval augmented generation (RAG) capabilities were recommended. In this study, we demonstrated with Qwen1.5, but we also provided codes compatible with GPT-4. Qwen1.5-72B-chat were downloaded from Hugging Face and deployed on a high-performance computing cluster with Nvidia A100 GPUs. The text from previous step were sequentially input into LLM with the following system prompt:

> *You are specialized for analyzing scientific paper abstracts, focusing on identifying entities causal relationships related to biological studies, such as performance, species, genes, methods of genetic engineering, enzymes, proteins, and bioprocess conditions (e*.*g*., *growth conditions). It outputs the identified causal relationships between entities in combination pairs. The output strictly follows the format: (Entity A, Entity B), (Entity C, Entity D)* … *with no additional text*.

The response from LLM was processed by the Word2Vec method ^24^. A word embedding from sentence-transformers ^25^ was used to identify and combine entities with same meanings. The processed lists of entities were saved and passed to the next step.

### Knowledge graph visualization

Each pair of entities indicates their causal relationship. An open-source Pyvis package ^26^ was used to construct knowledge graph. The nodes represent entities, while edges (lines connecting nodes) are labeled as the article title as citation/source.

### Keyword search and summarization

After obtaining the knowledge graph, it is recommended to search for a keyword and filter the knowledge graph for cleaner visualization. The entities/nodes related to the search keyword were extracted and input into LLM for summary report generation. LLMs with good reasoning ability, such as GPT-4 and Qwen1.5-32B/72B, are recommended.

### Problem solving and hypothesis generations

After obtaining summary reports about relevant concepts, these reports were combined. NEKO leverages the inference capabilities of LLMs to propose novel research plans and hypotheses. In the cyanobacterial nitrogen fixation example, concepts of *Anabaena* sp. nitrogen fixation and common bio-fertilizer formulation were input to Qwen1.5-72B. The LLM inference gives a brief description of potential research directions and hypotheses, which are evaluated by human researchers.

## Acknowledgements

This study is funded by United States NSF award number 2225809 and DOE Energy Earthshots award number DE-SC0024702.

## Author contributions

Conceptualization and Ideas - YJT, YC, ZX; Programming – GPT-4, Qwen, ZX; Writing – GPT-4, Qwen, ZX; Review & Editing -YJT, YC, HBP

## Supplementary files

Supplementary file 1: Supplementary Figures

