## supplementary file 1 for "Network for Knowledge Organization (NEKO): an AI knowledge mining workflow for synthetic biology research"

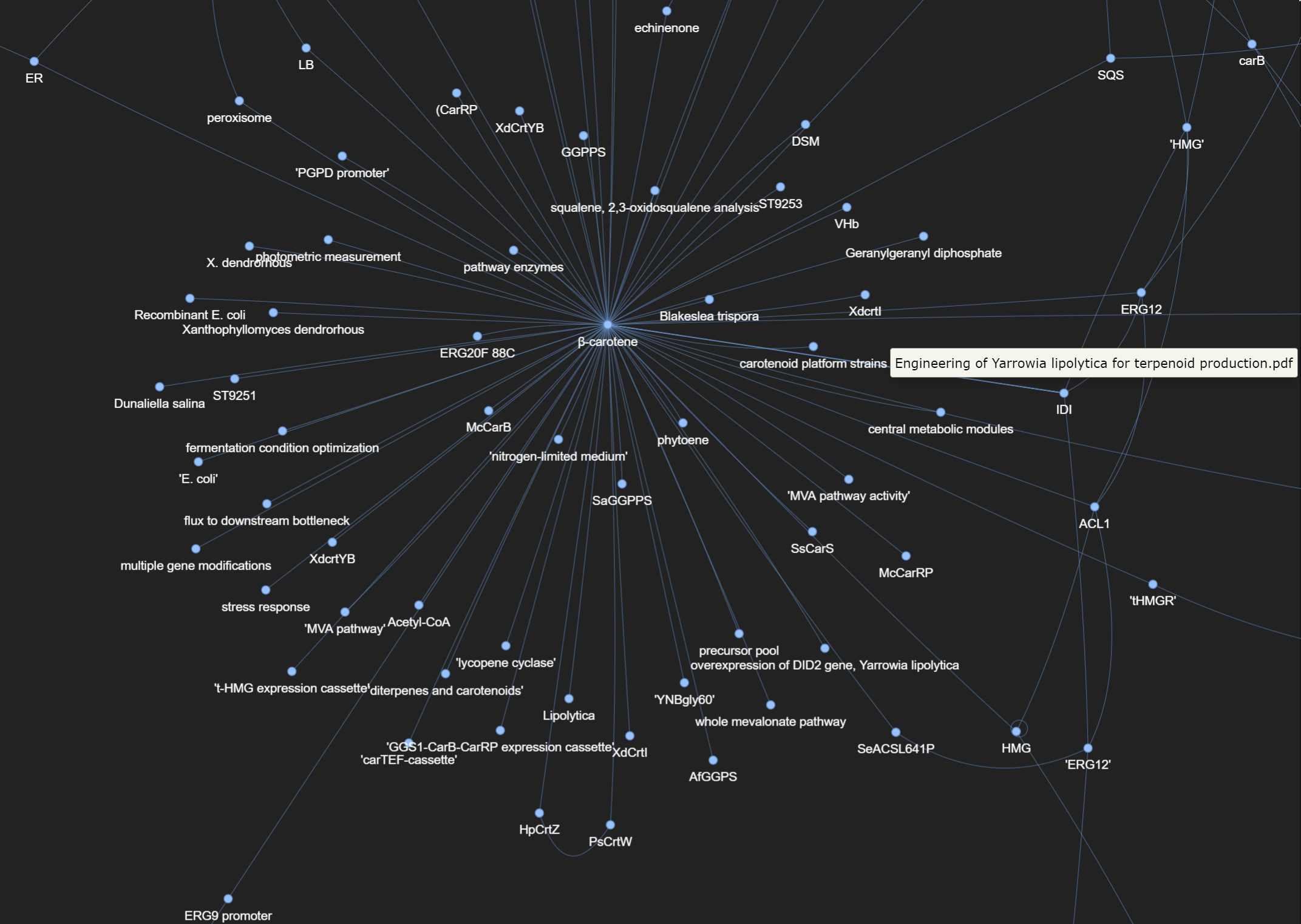

**Figure S1.** Part of β-carotene production knowledge graph. User can click a line connecting two nodes and view the title of the article containing the knowledge connection. The user can also search for knowledge citation in the article/entity list.

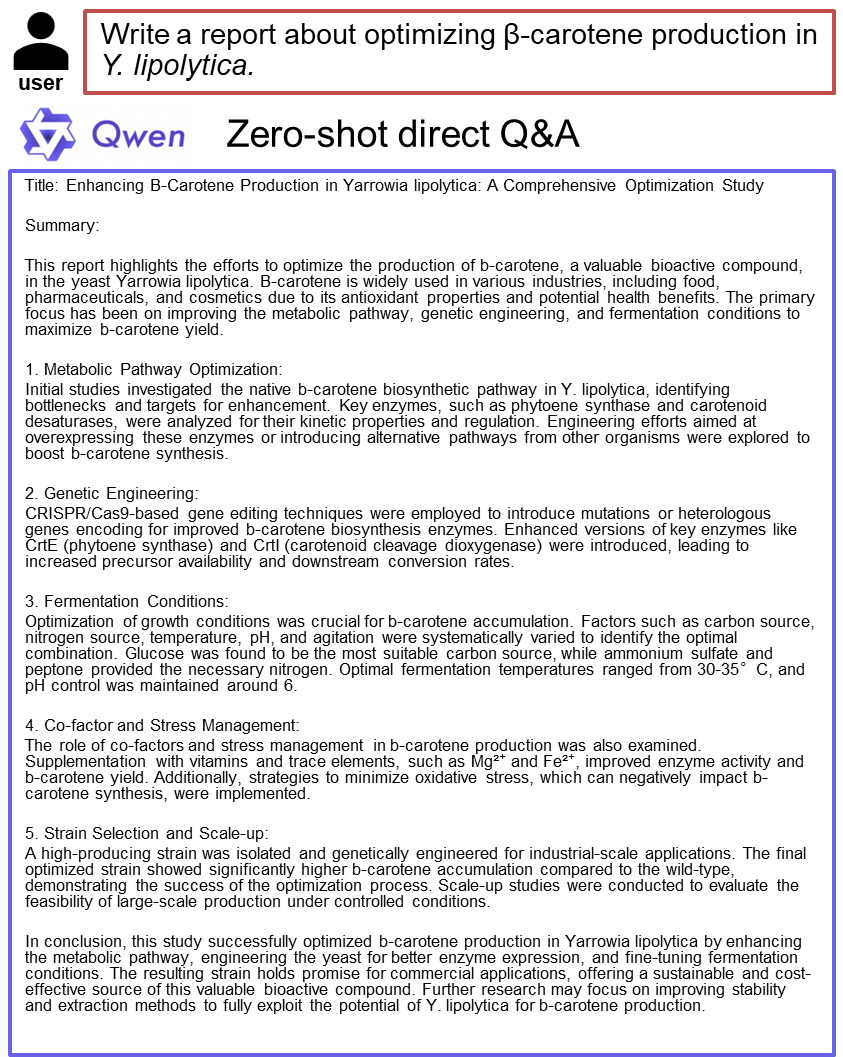

**Figure S2.** Qwen zero-shot Q&A summary of β-carotene production in Y. lipolytica. The base LLM Qwen does not have prior knowledge of β-carotene production. Only with NEKO workflow, LLM can output specific knowledge.

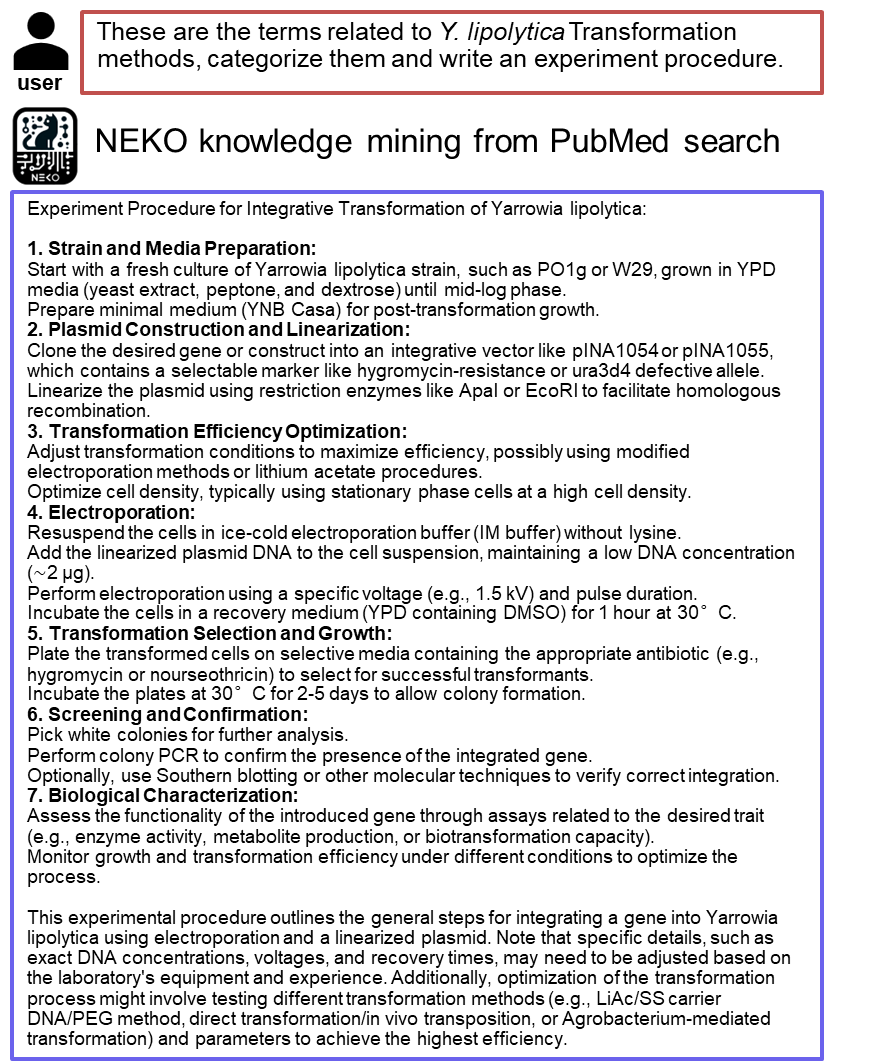

**Figure S3.** NEKO summarizes *Y. lipolytica* strain transformation procedures.

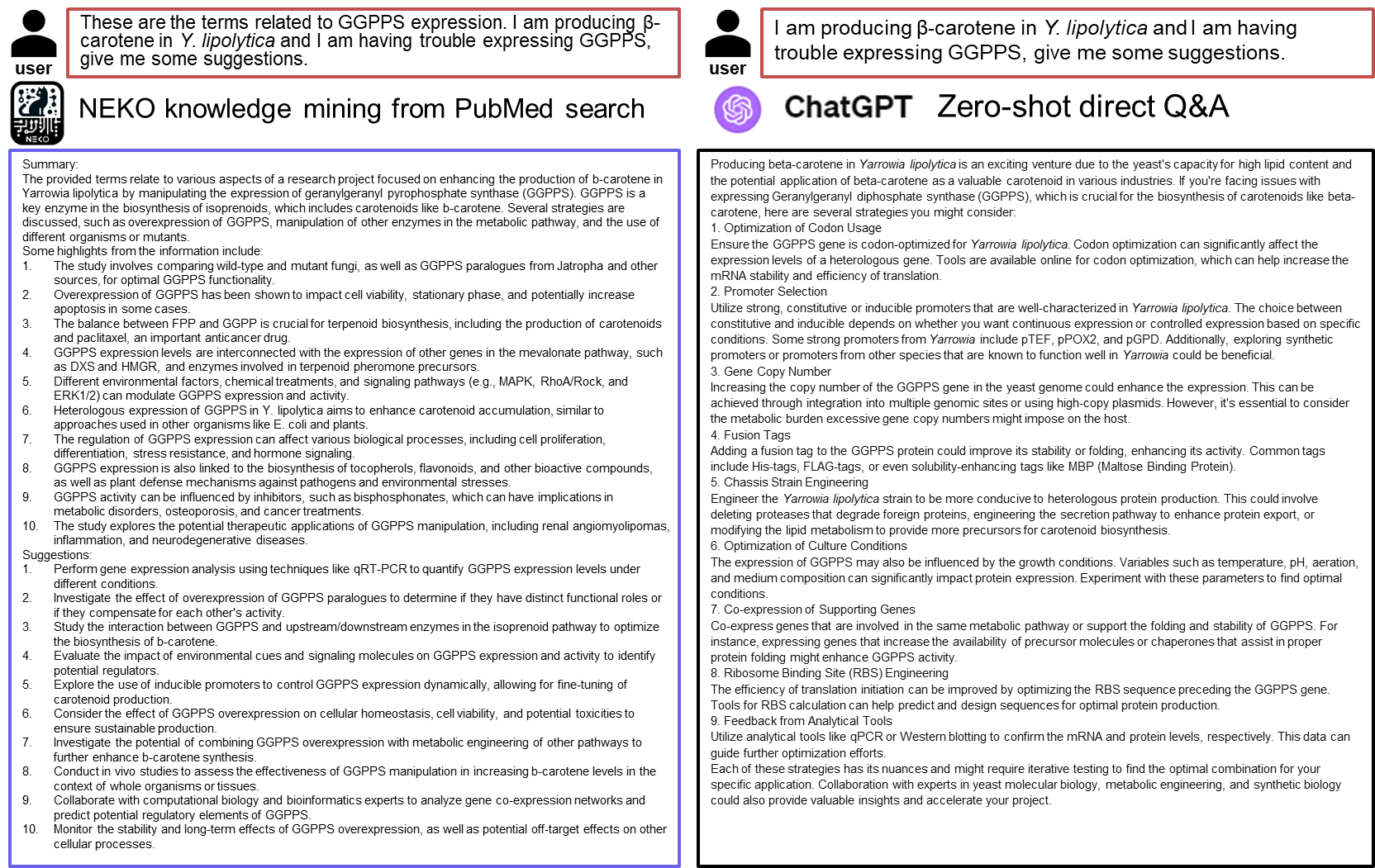

**Figure S4.** Compare NEKO and ChatGPT zero-shot Q&A about troubleshooting GGPPS expression.

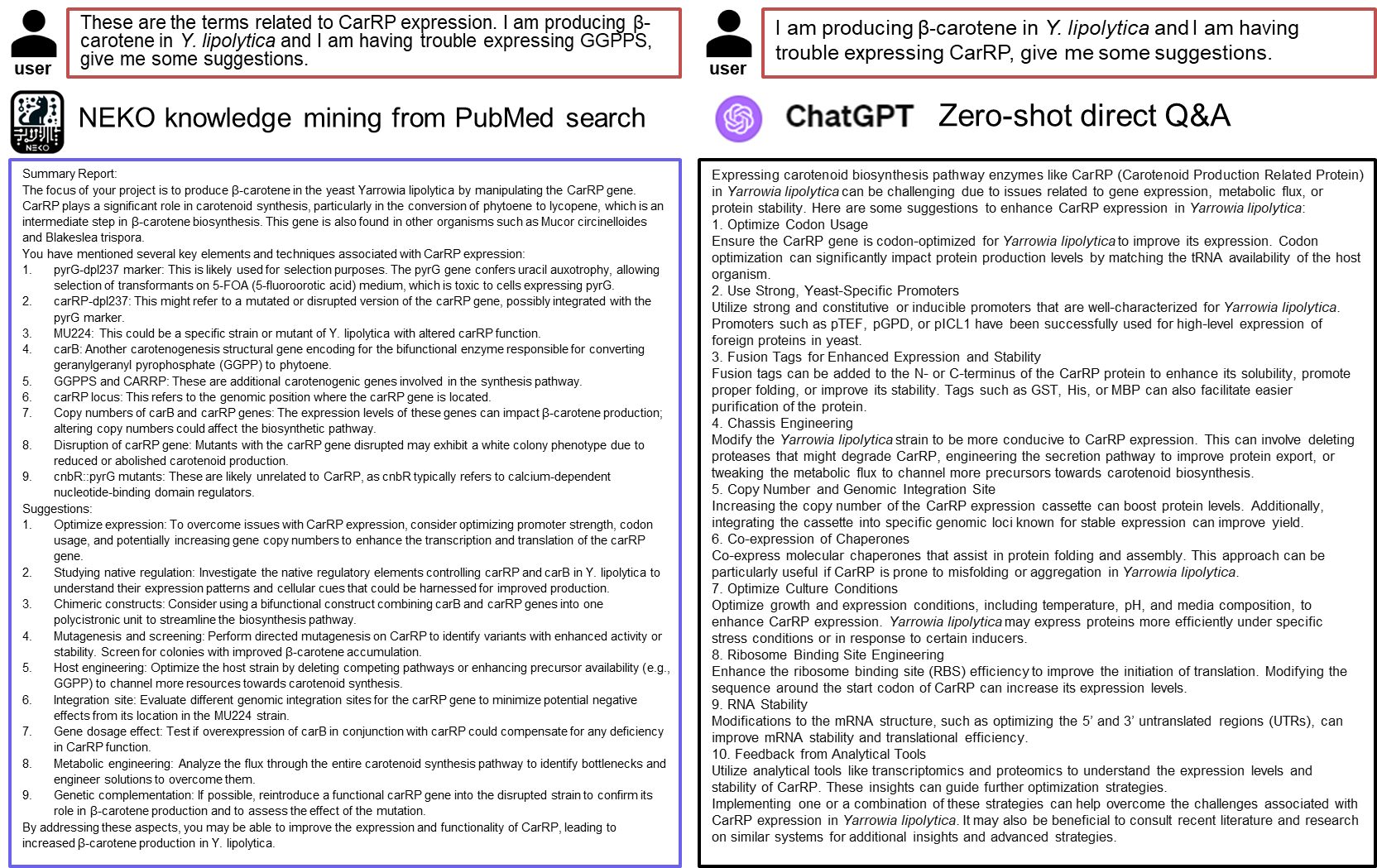

**Figure S5.** Compare NEKO and ChatGPT zero-shot Q&A about troubleshooting CarRP expression.

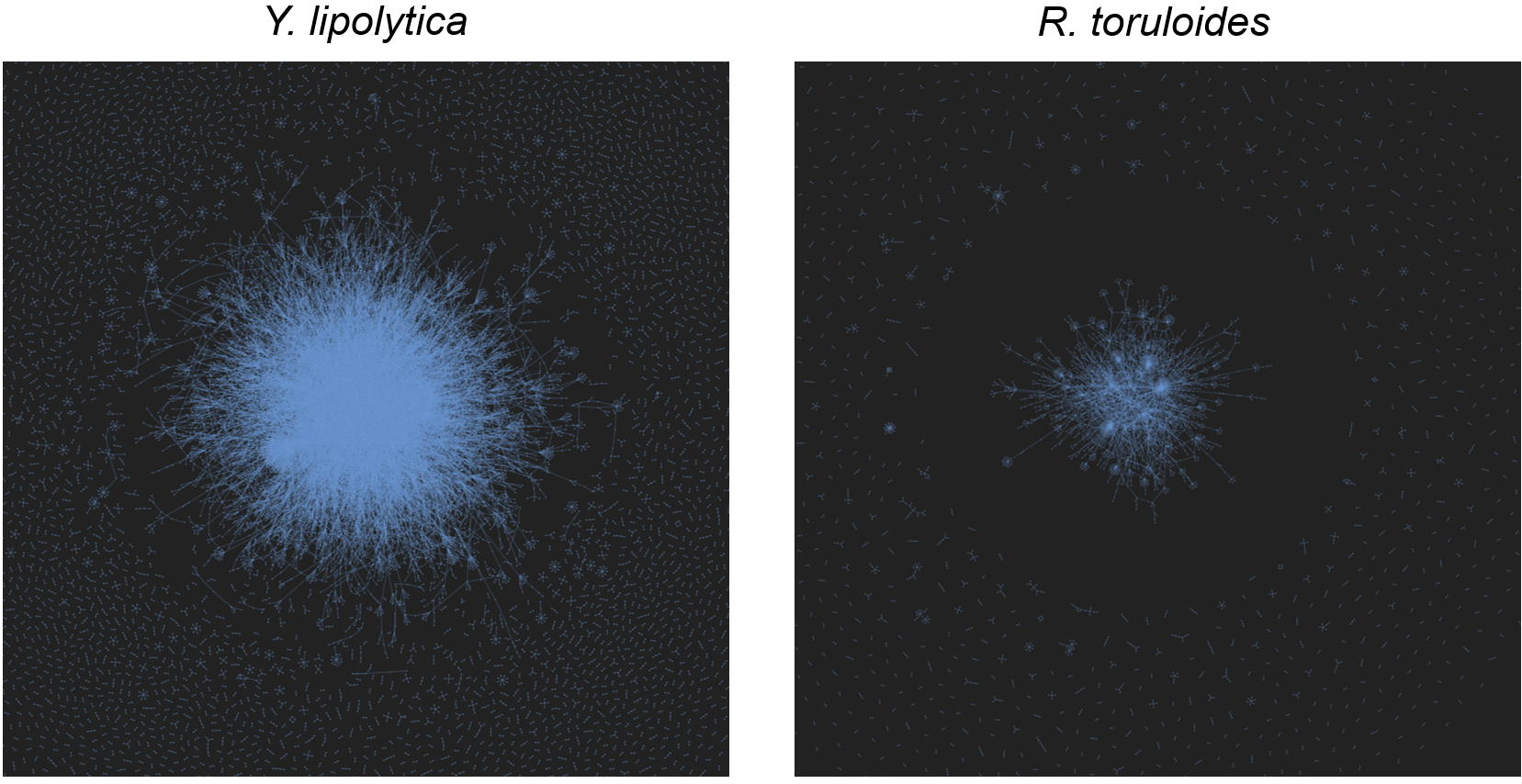

**Figure S6.** Compare knowledge graphs of *Y. lipolytica* and *R. toruloides*. The more studied *Y. lipolytica* has more connections in the core knowledge region.

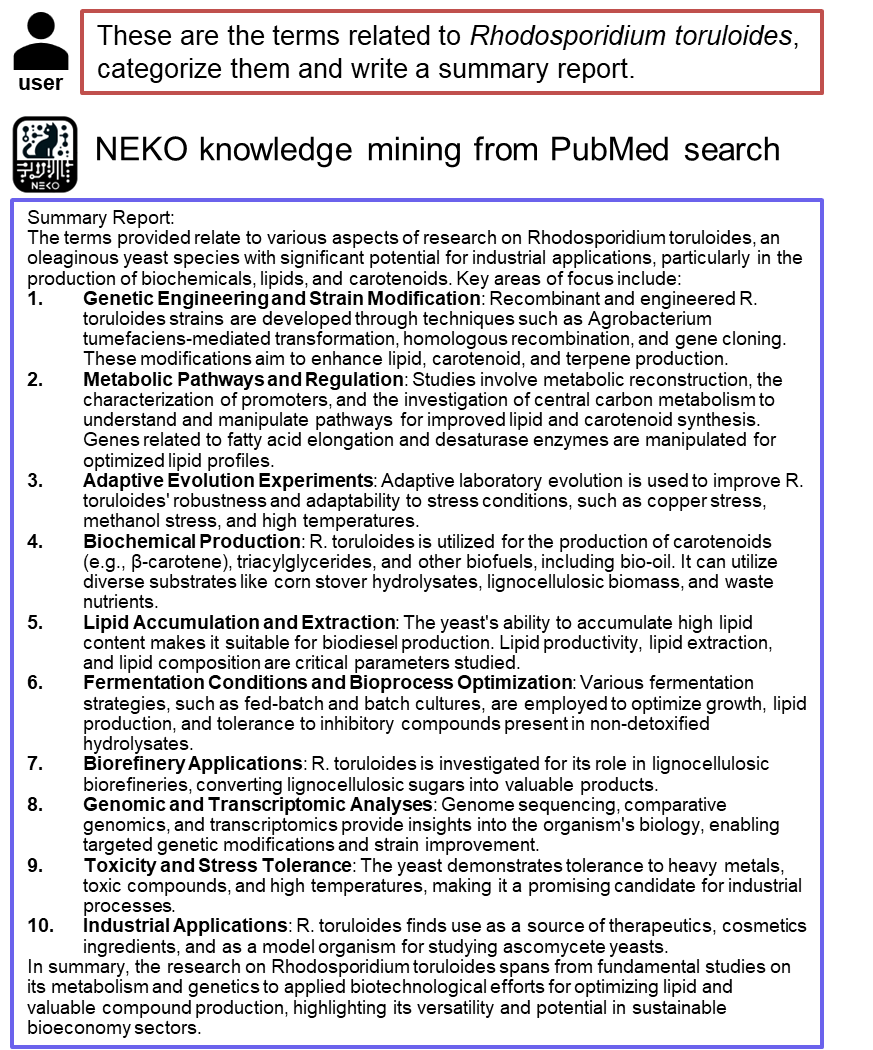

**Figure S7.** NEKO’s literature review on a non-model species *Rhodosporidium toruloides*. It identified trending research topics in literature.

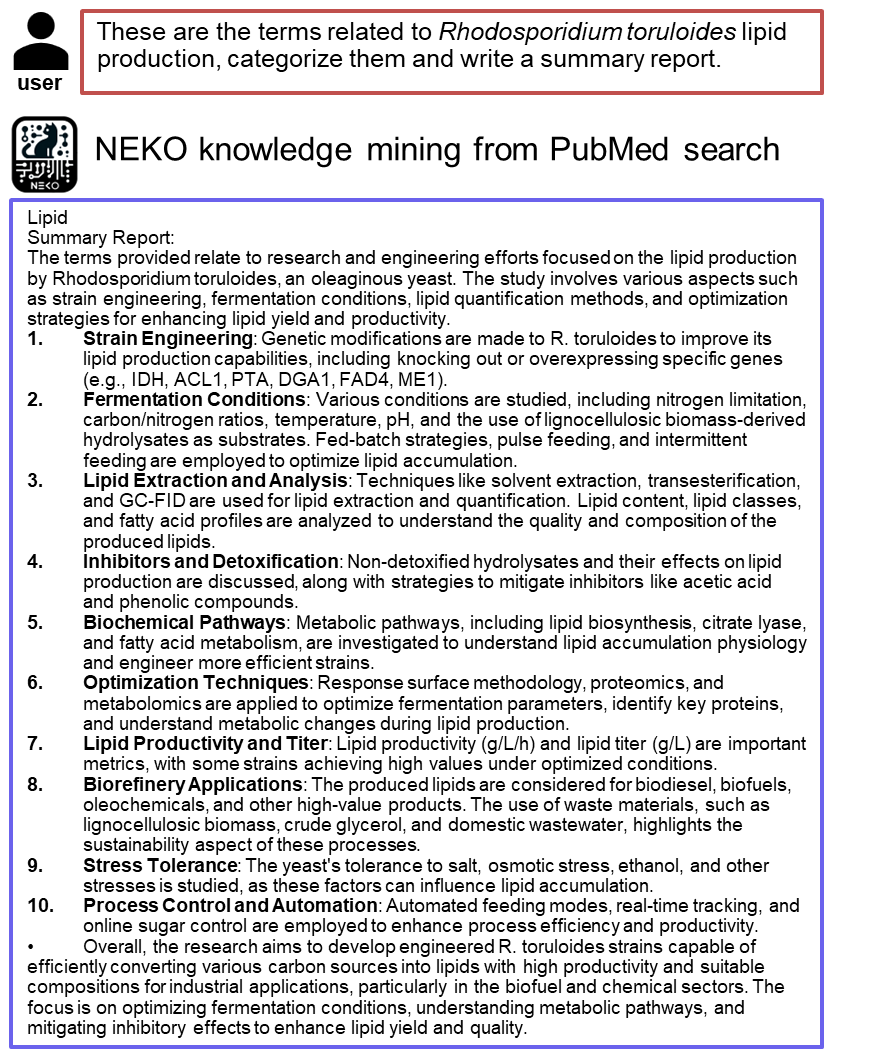

**Figure S8.** NEKO summarizes lipid production studies in *Rhodosporidium toruloides*.

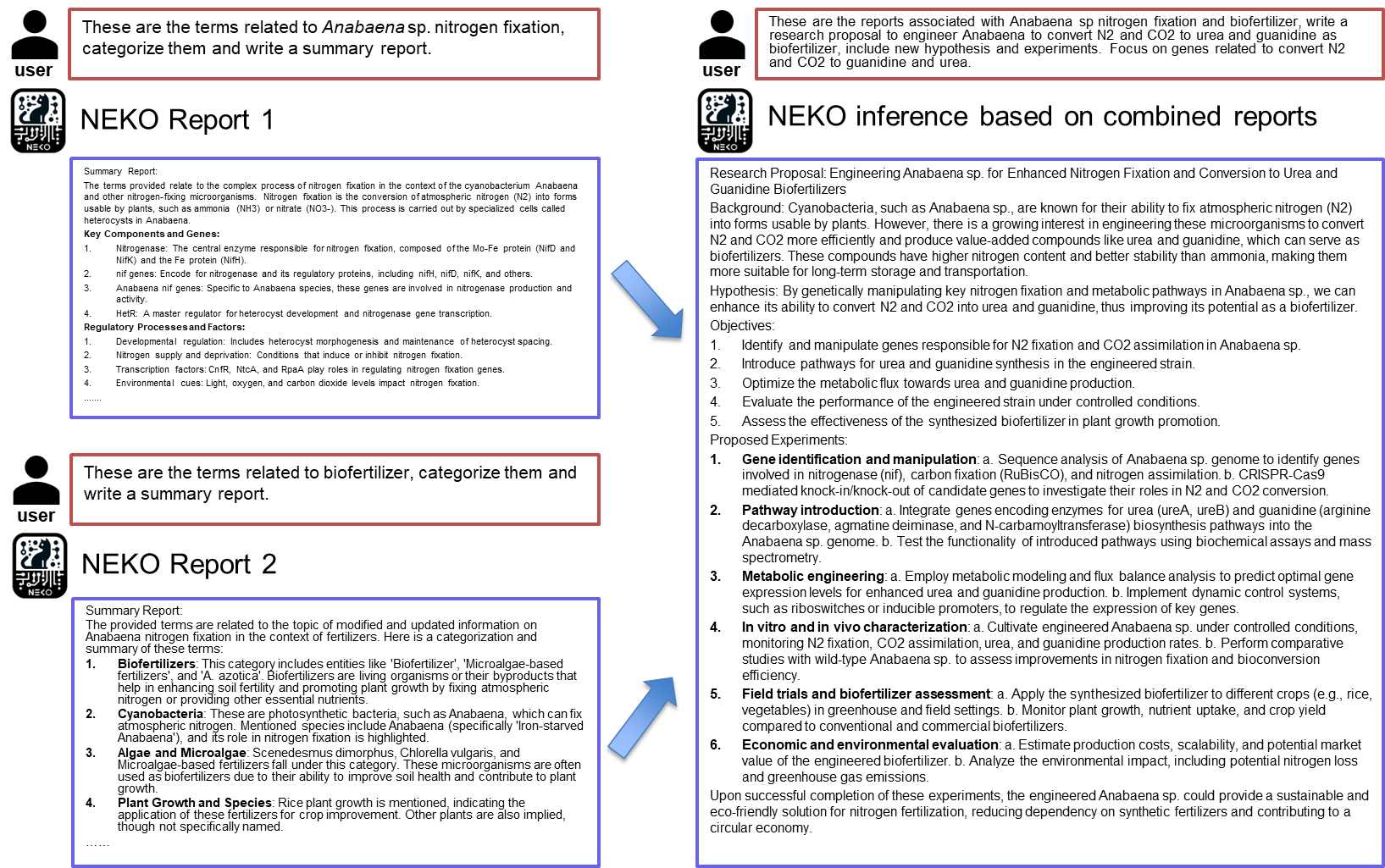

**Figure S9.** NEKO combines literature review reports. LLM inferences can identify potential research objectives.

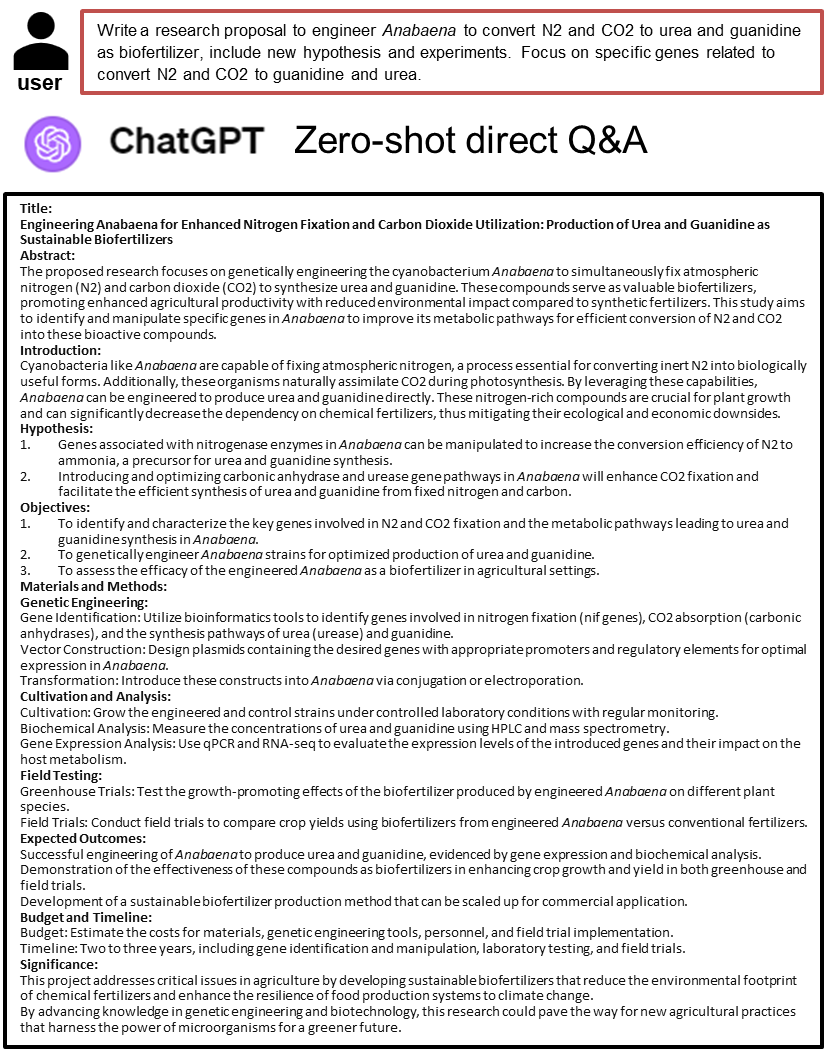

**Figure S10.** ChatGPT zero-shot direct Q&A about engineering *Anabaena* sp. for biofertilizer.

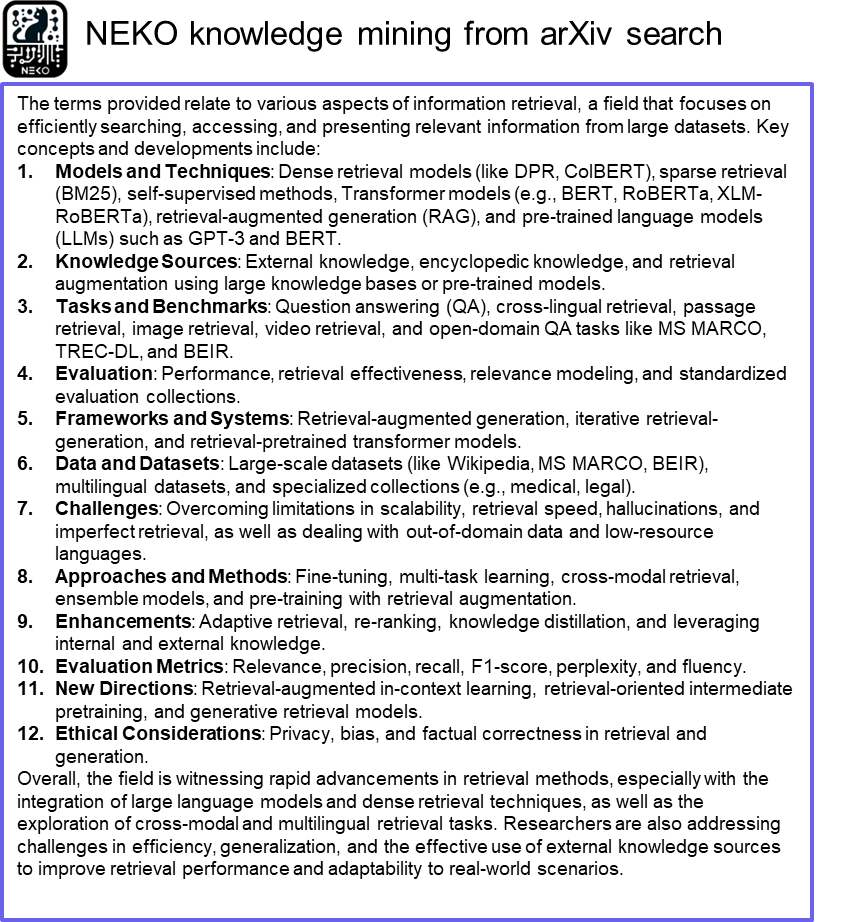

**Figure S11.** NEKO summarizes LLM knowledge retrieval from arXiv computer science articles.
